# A Mouse-adapted SARS-CoV-2 Model for Investigating Post-acute Sequelae of COVID infection

**DOI:** 10.1101/2024.11.10.622868

**Authors:** Haowen Qiao, Yafei Qu, Lingxi Qiu, Yuanpu Chiu, Xiao He, Tenghuan Ge, Zhen Zhao, Weiming Yuan

## Abstract

The coronavirus disease of 2019 (COVID-19), caused by the Severe Acute Respiratory Syndrome Coronavirus-2 (SARS-CoV-2), remains a major health issue after nearly 7 millions of death toll in the last four years. As the world is recovering with improving vaccines and antiviral treatments, the alarming rate of long-COVID, or Post-acute Sequelae of COVID-19 (PASC), calls for further investigations. Among a list of symptoms associated with multi-organ dysfunctions, the neurological complications are particularly intriguing, yet the underlying mechanisms remain elusive. With the recently developed mouse adapted SARS-CoV-2 stain, we are now able to model the mild COVID infection in C57BL/6 mice and study the chronic immune responses and subsequent damages in different organs long after the viruses are clearly naturally in the body. More specifically, we found adult C57BL/6J mice developed neurological impairments, including behavior changes related to sensorimotor coordination, depression- and anxiety-like behaviors, and inflammation in multiple organs including lung, liver and brain, which persisted over at least 4 weeks in mice even with mild infection. Therefore, this model can be used to further explopred the mechanisms of PASC, as well as potential intervention or therapeutic approaches.

## Introduction

After several devastating waves of worldwide epidemics, although the mortality and virulence of SARS-CoV-2 have substantially dropped, the extended post-acute sequelae of COVID-19 (PASC, also known as long COVID) still represents a major challenge of public health for millions of patients^1,2^. Long COVID is a multi-organ disease involving dysfunctioned or dysregulated immune, respiratory, cardiovascular, gastrointestinal, neuropsychological, musculoskeletal, and/or other systems, which persist at least 4 weeks after the exposure to SARS-CoV-2^3,4^ and may last life long. Among the multi system sequelae in long COVID, neurological complications are most common and are highly challenging to manage^4^. Retrospective studies revealed that neurological manifestations were prevalent in hospitalized COVID-19 patients^5^. Stroke, venous thromboembolism, intraventricular and subarachnoid hemorrhage, encephalitis and seizures were observed in more than 30% of the patients^6–10^. More importantly, neurological PASC were also frequently reported in COVID patients who only experienced with mild infection^11–17^. For example, the RECOVER adult cohort study that among the 37 symptoms, postexertional malaise (PEM), fatigue, dizziness and brain fog are most frequent^13^. Therefore, it remains an unmet need to understand the neuroinvasiveness of SARS-COV-2 virus and its impact on the central nervous system at fundamental levels^18–23^.

Both rodents and nonhuman primates have been used host systems to study SARS-COV-2 virus infection, and long COVID. Although mice are not natural host of SARS-CoV-2, this issue was first overcome by the transgenic models carrying the human ACE2 receptor^24^, and more recently by reverse engineering the viral genome and/or serial passaging *in vivo*^25–27^. The newly developed mouse adapted SARS-CoV-2 strains have largely mitigated the resistance issues, and are extensively characterized based on their *in vivo* passages and virulence^28^. For example, the MA10 strain carries amino acid changes Q498Y and P499T of spike protein, and enables viral entry through the murine ACE2 receptor^25,28^. More importantly, the adapted strains usually elicit mild symptoms, and can recapitulate key manifestations of the PASC including neuropathologies^25,28–34^. Furthermore, mouse adapted strains are easy to implement on mice with different genetic background (e.g., transgenic or knockout models), compared to other alternative models such as AAV-mediated restricted expression of hACE2 in lung^31^ or humanized mouse model (e.g. MISTRG6 model with transplantation of human hematopoietic stem and progenitor cells)^34^.

To study the mechanisms of neurological sequelae caused by SARS-CoV-2, we established a murine model of long COVID diseases in C57BL/6J mice with the well-established SARS-COV-2 MA10vstrain^25^. After thorough surveillance and validations of viral load and shedding, mice no longer carry active virus at 14 days after inoculation were transported from the animal biosafety level 3 (ABSL-3) facility, allowing detailed analyses of mouse behavior over time, followed by neuropathological analysis. We found that even with mild infection with SARS-CoV-2 MA10 virus, adult C57BL/6J developed neurological impairments, including behavior changes related to sensorimotor coordination, depression and anxiety-like behaviors based on different assessments in 2-4 weeks after initial infection. More importantly, histological analyses revealed that neuroinflammation occurred in the brain after SARS-CoV-2 MA10 infection and viral clearance in the body, which even persisted longer than the changes in peripheral organs, such as lung and liver. Our findings demonstrated that mild infection model in C57BL/6J mice with SARS-CoV-2 MA10 strain can be used to understand the pathogenesis and progression of PASC.

## Results

### Behavior changes in mice after SARS-CoV-2 MA10 infection

10-12 weeks old C57BL/6J mice (male and female) were infected with MA10 virus, and we found that different infectious dose or regimen caused different outcomes. C57BL/6J mice infected at 6×10^4^ pfu/mouse in 50 ul of PBS exhibited minimal weight loss without lethality (**Figure 1a**), while higher dose can cause significant weight loss in first 7 days after infection, sometime even surpass the 20% weight loss limit where animals had to be euthanized. Therefore, we considered mice with weight loss was below 10% (compared with the initial weight) as mild infection, while mice with weight loss greater than 10% were defined as moderate, and euthanized mice wer considered as severe infection (**Figure 1a**).

**Figure 1.**
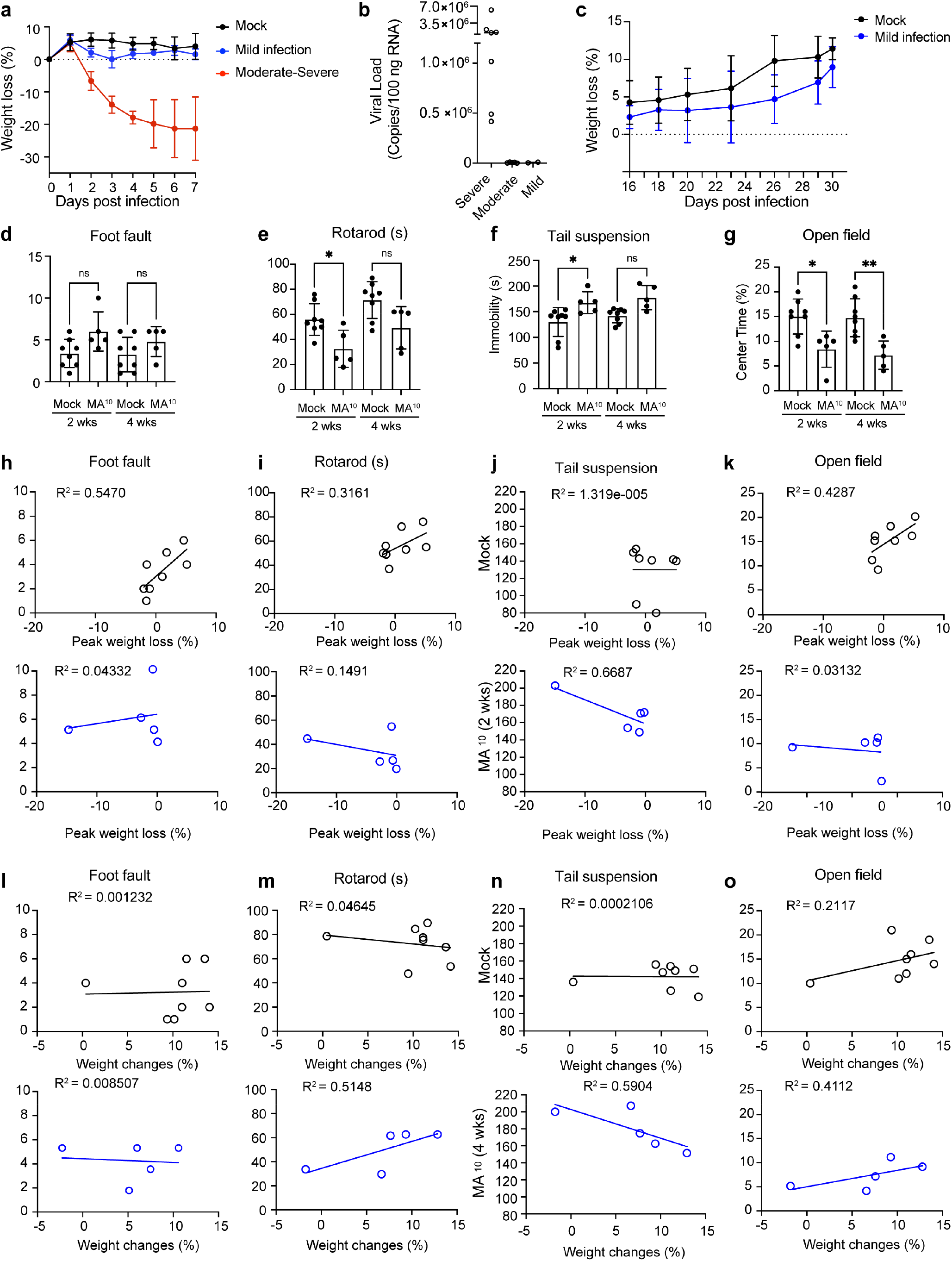
SARS-CoV-2 MA10 infection modified the behavior of mice. **a**. Body weight changes (%) of mock (n=5) and MA10-infected (n=5) C57BL/6J mice during the first week post inoculation. Body weight up to 20% in moderate-severe infection conditions were plot in red as a comparison. **b**. Viral copies of MA10 in 100 ng total RNA determined by qPCR of N1 transcript (see methods). 14 mice from mild infection (n=2), moderate (n=5) or severe (n=7) infection were analyzed. **c**. Body weight changes (%) of mock (n=4) and MA10-infected (n=4) C57BL/6J mice measured in ABSL-2 facilty. **d-g**. Behavioral tests for mock (n=8) and MA10-infected (n=5) C57BL/6J mice at 2- and 4-weeks post inoculation, including foot fault test (**d**), rotarod test (**e**), tail suspension test (**f**) and open field test (**g**). mean ± SD; *, p < 0.05; **, p < 0.01; ns, not significant, by two-tailed unpaired Student’s t-test. (**h-k**) Linear correlation analyses of mock (upper panel, n=8) or MA10-infected (lower panel, n=5) C57BL/6J mice between peak weight loss (%) and their behavior performance at 2 weeks post inoculation. R^2^, coefficient of determination of the linear relationship. (**l-o**) Linear correlation analyses of mock (upper panel, n=8) or MA10-infected (lower panel, n=5) C57BL/6J mice between final weight changes (%) and their behavior performance at 4 weeks post inoculation.

Tissue samples were collected from a corhort of mice with mild to severe conditions, and homogenized in TRIzol reagent for RNA extraction. The complementary DNA (cDNA) were used for qRT-PCR analysis of viral RNA copies (see Methods). Residual low-copies of viral RNA molecules can still be detected in the lungs from mice with mild to moderate infections at dpi 14 (**Figure 1b**), compared to the high copy numbers detected in severely infected mice. This is consistent with reports from patients that persistent vRNA detection after 2 years in convalescent patients^35^. More importantly, viral shredding was negative based on quantitative RT-PCR analysis from body wipes or feces samples, indicating a natural clearance in mice as observed in patients.

For the current study on long COVID / PASC, we focus on the mild infection model. Once the mice were validated to be free of live virus, they were transferred to ABSL-2 for maintenance and behavioral testing. The mild infected mice gradually gained weight over time (**Figure 1c**) and showed no significant difference in activity from the mock group. At 15 and 30 dpi, we performed an array of behavioral tests to examine the neurological sequelae. We evaluated sensorimotor coordination with foot fault and rotarod tests (**Figure 1d-e**), depression-like behavior with tail suspension test (**Figure 1f**) and anxiety-like behavior with open field test (**Figure 1g**). The data were collected at 15 and 30 dpi, and summaried in **Table 1**. Interestingly, we observed that the convalescent mice demonstrated reduced sensorimotor functions temporarily at 15 dpi, based on a slight increase in foot fault numbers (**Figure 1d**) and significant reduction in riding time on rotarod test (**Figure 1e**). On the other hand cognitive changes lasted longer, as an increase in depressive behavior based on immobility time on Tail suspension test (**Figure 1f**) and a significant reduction in time spent in the center area during the Open field test (**Figure 1g**) were still observed at 30 dpi.

**Table 1.**
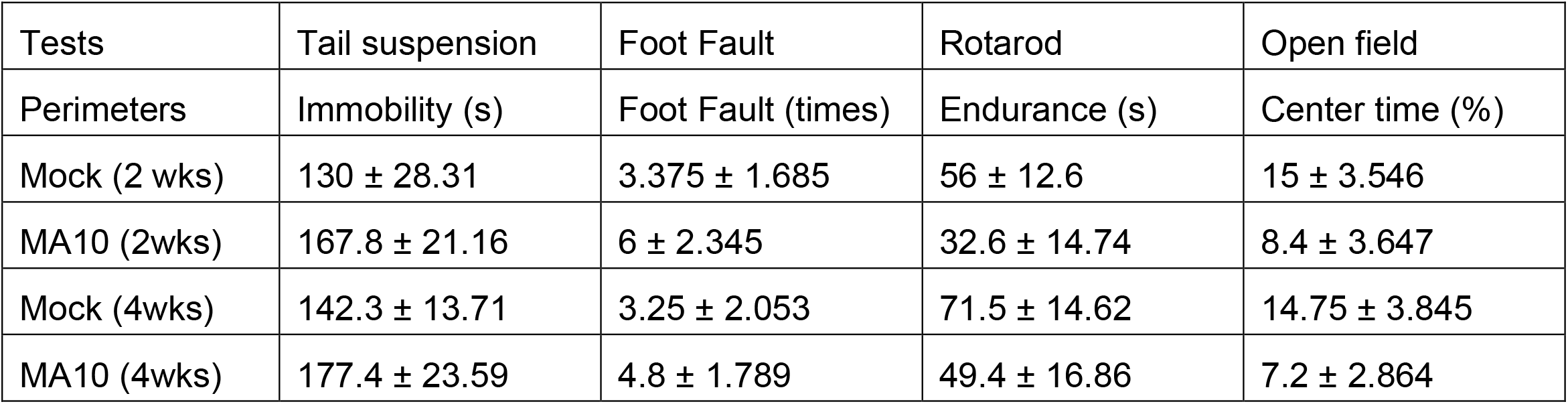
The statistical analysis of behavioral data.

| Tests | Tail suspension | Foot Fault | Rotarod | Open field |
| --- | --- | --- | --- | --- |
| Perimeters | Immobility (s) | Foot Fault (times) | Endurance (s) | Center time (%) |
| Mock (2 wks) | 130 ± 28.31 | 3.375 ± 1.685 | 56 ± 12.6 | 15 ± 3.546 |
| MA10 (2wks) | 167.8 ± 21.16 | 6 ± 2.345 | 32.6 ± 14.74 | 8.4 ± 3.647 |
| Mock (4wks) | 142.3 ± 13.71 | 3.25 ± 2.053 | 71.5 ± 14.62 | 14.75 ± 3.845 |
| MA10 (4wks) | 177.4 ± 23.59 | 4.8 ± 1.789 | 49.4 ± 16.86 | 7.2 ± 2.864 |

Based on the fact that SARS-CoV-2 MA10 infection changed the behavior of mice, we studied the correlation between weight changes and behavioral changes (**Figure 1h-o**). The body weight loss caused by viral infection usually peaked within one week post infection, and then infected mice gradually regained their weight. We first analyzed the relationship between peak weight loss and the behavioral outcomes at 15 dpi (**Figure 1h-k**). Interestingly, the peak weight loss was only related to behavior changes in Tail suspension test (**Figure 1j**), but no significant correlation with motor coordination (**Figure 1h-i**) or anxiety-like behavior (**Figure 1k**) was observed. As mouse body weight fully returned to or even surpass their initial weight at four weeks post infection, we next studied the correlation between final weight changes and behavior outcomes at 30 dpi (**Figure 1l-o**), and found that the final weights of infected mice had a correlation with behavior changes from both rotarod test (**Figure 1m**) and Tail suspension test (**Figure 1n**), and marginally correlated with anxiety-like behavior from Open field test (**Figure 1o**). Therefore, our data indicated that mild infection with SARS-CoV-2 MA10 can cause PASC in mice.

### SARS-CoV-2 MA10 infection induced chronic inflammation in the lung and liver

Mast cells activation symptoms were reported to be prevalent in patient with PASC^36^. Tissue histology with Toluidine Blue staining revealed the presence of granulated mast cells in the lung of SARS-CoV-2 MA10 infected mice at 3 dpi, which was resolved later at 14 and 30 dpi (**Figure 2a-b**). Additionally, we observed accumulation of CD68+ macrophages near the bronchioles at 3 dpi in response to SARS-CoV-2MA infection, which gradually returned to normal from 14 to 30 dpi (**Figure 2c-d**). We also assessed macrophage dynamics in the liver over time, and only found a temporal increase of CD68+ macrophages at 14 dpi following SARS-CoV-2 infection (**Figures 2e-f**).

**Figure 2.**
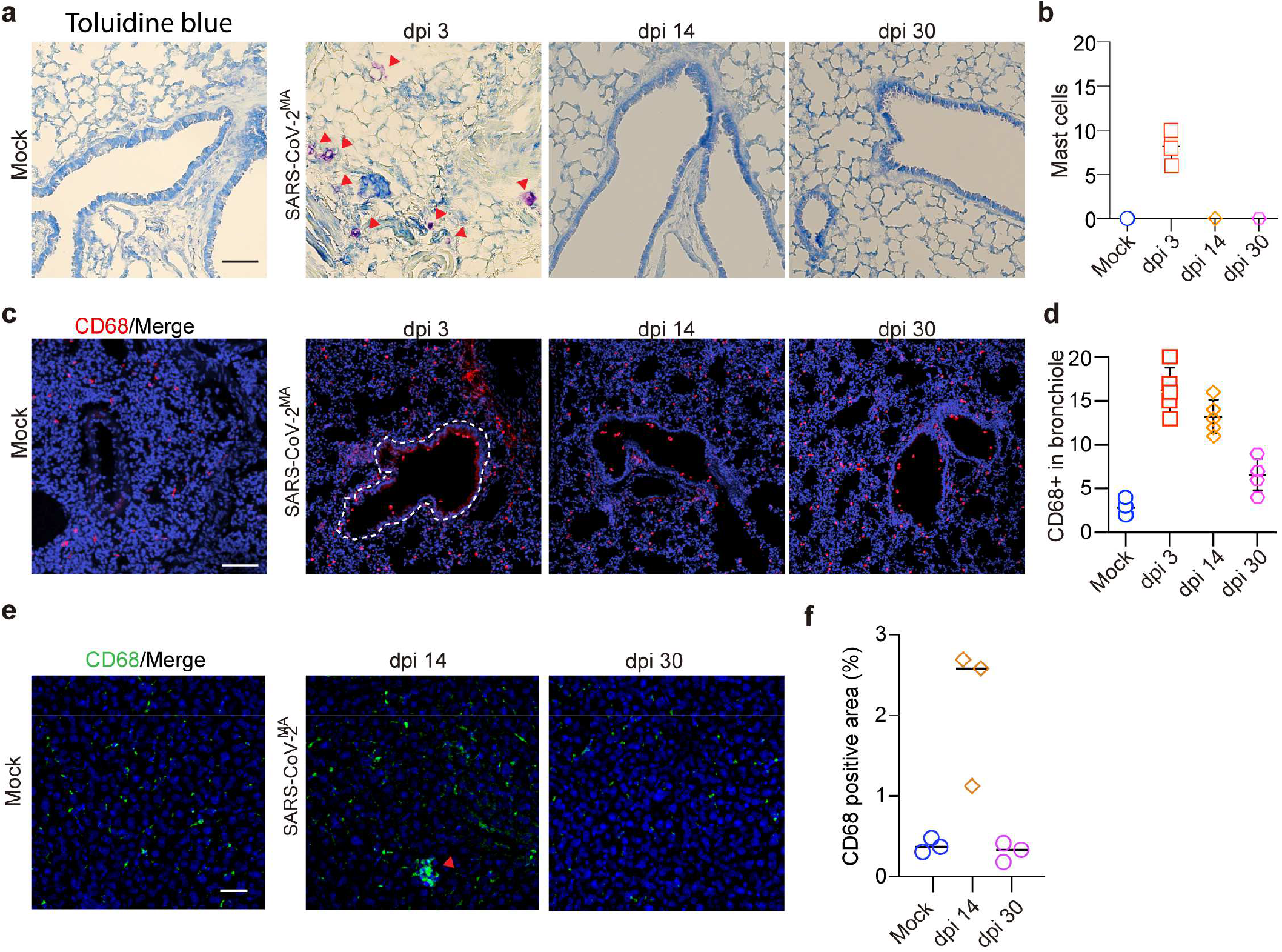
SARS-CoV-2 MA10 infection induced inflammation in the lung and liver. **a**. Histological images of toluidine blue-stained trachea sections from Mock or SARS-CoV-2 MA10 infected C57BL/6J mice. Mast cells (arrowheads) are observed in the trachea of SARS-CoV-2-infected mice at 3 dpi. **b**. Quantification of mast cell counts at 3, 14, and 30 dpi from Mock or SARS-CoV-2 MA10 infected mice. n=3 mice per group. **c**. Representative images showing immunofluorescence staining of CD68-postive macrophages in the lung of Mock or SARS-CoV-2 MA10 infected mice. **d**. Quantification of CD68+ macrophage counts in the bronchiole area. n=3-5 mice per group. **e**. Representative images showing immunostaining of CD68-postive macrophages in the liver. Arrowhead indicates a cluster of active macrophage, which is only observed in SARS-CoV-2 MA10 infected mice (dpi 14). **f**. Quantification of CD68+ macrophage positive area in the liver. n=3 mice per group.

### SARS-CoV-2 MA10 infection triggered neuroinflammation in brain

Choroid plexus is a known target of COVID infection based on recently single cell transcriptomics study^37^, therefore, we next examined the border-associated macrophages (BAM) in the brain of SARS-CoV-2 MA10 infected mice. These BAMs are capable of leaving the choroid plexus, reaching the ventricular wall and breaching the ependymal cells after infection^38,39^ (**Figure 3a**). We found that SARS-CoV-2 MA10 infection led to an significant accumulation of Iba1+ macrophages in the choroid plexus (**Figure 3b-c, Supplementary Figure 1a-b**), with some of the Iba1+ cells localizing to the apical, cerebrospinal fluid (CSF)-facing surface of the ChP (**Figure 3d, Supplementary Figure 1c**), which is quite different from mock condition where most of the Iba1+ cells remains within the choroid plexus. To further define the identity of the BAM macrophages lining the wall, we immunostained for BAM marker CD206, and found SARS-CoV-2 MA10 infection resulted in an dramatic increase in CD206+ BAMs both on the ventricular wall and inside the ependymal cell layer (**Figure 3e-h**). This observation was further confirmed in both lateral and third ventricles (**Supplementary Figure 1d-h**).

**Figure 3.**
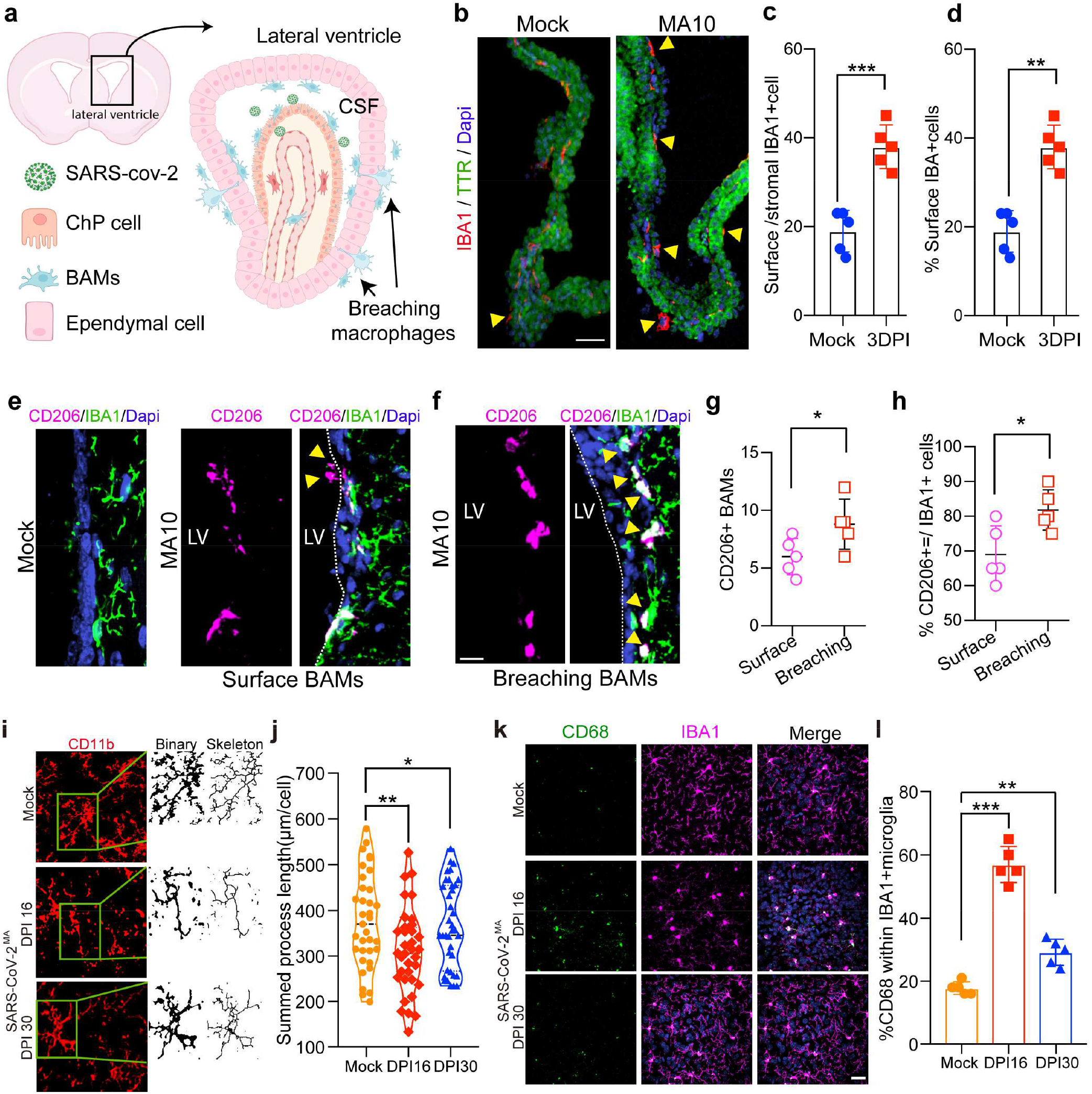
SARS-CoV-2 MA10 infection triggered neuroinflammation in the brain. **a**. Schematic illustrating the process of extracting border macrophages (BAMs) that breach the lateral ventricle (LV) lining after SARS-CoV-2 infection in mice. **b**. Representative images showing choroid plexus (ChP) of LV with IBA1 macrophages from mice with or without SARS-CoV-2 MA10 infection. TTR, transthyretin, marker of ChP. Arrowheads indicate migratory macrophages located at the ChP surface. Scale bar: 50 µm. **c**. Proportion of surface ChP macrophages relative to stromal macrophages. Data from n = 5 mice; ***p < 0.001, determined by two-tailed Student’s t-test. **d**. Percentage of macrophages found on the surface of ChP. Data from n=5 mice; **p < 0.01, determined by two-tailed Student’s t-test. **e-f**. Representative images demonstrating CD206-positive BAMs near the surface of the ependymal cell layer (**e**) or breaching into the ependymal cell layer, following SARS-CoV-2 MA10 infection. **g**. Quantification of CD206+ BAM counts at the surface of the ventricles, and breaching into the ependymal cell layer. **h**. Quantification of the percentage of CD206+ BAMs among IBA1+ cells at the surface of the ventricles, and breaching into the ependymal cells. **i**. Representative images showing immunostaining of CD11b-positive microglia in cortex region of mice with or without SARS-CoV-2 MA10 infection. Binary and skeletonized images were produced using bandpass filter and the Skeleton plugin in ImageJ, respectively. **j**. Dot-plot graphs showing total process length per microglia analyzed (n = 35 cells from 4 mice in each group). SARS-CoV-2 MA10 infected mice were examined at 16 and 30 days post infection (DPI). **k**. Representative images showing immunostaining staining of CD68 and IBA1 marker in the cortex from SARS-CoV-2MA-infected C57BL/6J mice. Bar = 25 μm. **l**. Quantifi-cation of percentage of CD68+ active microglia in IBA1+ total microglia. n = 5 mice per group. **p < 0.01; ***p < 0.001; one-way ANOVA with Bonferroni’s post hoc test.

To explore the long-term effects of infection, we followed the mice for 4 weeks and assessed morphological changes in cortical microglia in mice with Mock or SARS-CoV-2 MA10 infection. We discovered that even mild infection resulted in microglia activation at both 16 and 30 dpi, based on microglia morphological analysis (**Figure 3i-j**).and activation marker CD68 (**Figure 3k-l**). Briefly, we converted confocal image of CD11b cells into binary with thresholding method and skeletonization plugin, and quantified the total length of microglia processes between different group of mice (**Figure 3i-j**), and found SARS-CoV-2 MA10 infection reduced the process length of microglia even at 30 dpi. We also compared the activated microglia that are CD68+ and IBA1+ double positive, and found significantly increased number of active microglia after SARS-CoV-2 MA10 infection, at both 16 and 30 dpi (**Figure 3k-l**). Taken together, these results indicate that SARS-CoV-2 MA10 mild infection model in C57BL/6J mice also recapitulates the persistent neuroinflammation phenotype.

## Conclusion

So far, only limited number studies were able to use mouse models to explore the in-depth mechanisms of long COVID or PASC, including K18-hACE2 with low dose of infection^40–42^, AAV-mediated restricted expression of hACE2 in lung^31^, or Balb/c mouse strain^43^. However, the limitations due to rodent’s natural resistance has been lifted by the mouse adapted strains. To our knowledge, this is the first report of a long Covid model in C57BL6 strain using a well-characterized mouse adapted strain. We believe the model will be readily explopred for broader investigation on neuropathology and multi-organ complications associated with PASC, and also applicable to disease comorbidity such as diabetes and dementia models.

## Matierials and Methods

### Virus, mouse strains and SARS-CoV-2 infection

The mouse-adapted SARS-CoV-2 virus (MA10) was obtained from Biodefense and Emerging Infections Research Resources Repository (BEI Resources, NR-55329), propagated in Vero-E6 cells (ATCC, CRL-1586) as previously described^44,45^. C57BL/6J mice were purchased from the Jackson Laboratory (JAX #000664). All mice were maintained at Animal Biosafety Level 3 (ABSL-3) facility at the University of Southern California (USC) and approved by the Institutional Biosafety Committee (IBC) and the Institutional Animal Care and Use Committee (IACUC) at USC.

SARS-CoV-2 infections were performed in the Hastings and Wright Laboratories certified as an Animal Biosafety-Level 3 (ABSL3) containment laboratory at USC, following the Standard Operating Procedure (SOPs) provided by the ABSL3 facility as we previously reported^44,45^. Briefly, 10-12 weeks old C57BL/6J J mice (male and female) were intranasally inoculated with 6×10^4^ PFU of MA10 in 50 μL PBS, or PBS alone, under anathesia. Body weight and well-being of the mice were monitored daily once or twice for 14 days.

For the long-COVID study, each cohort of mice were validated to be free of live SARS-CoV-2 virus after 14 days by quantitative realtime PCR and plaque assays, and then transferred to from ABSL-3 to ABSL-2 holding with the approval from both IBC and IACUC. More specifically, viral shedding was examined by real-time PCR analysis of viral RNA copies in body wipes and feces samples at multiple time points between 3 to 14 days post-infection. In addition, one mouse from each cage was euthanized by isoflurane overdose followed by cervical dislocation, for more accurate analysis of viral in the lung by real-time PCR. Briefly, lung tissues were collected and homogenized in TRIzol reagent. Total RNA was extracted by Quick-RNA Miniprep Kit (Zymo Research, R1055) and used for RT-PCR (Bio-Rad, 1708840). The complementary DNA (dDNA) was used for qPCR analysis using Bio-Rad CFX Connect 96-well Real-Time qPCR module system with PrimeTime Gene Expression Master Mix (IDT, #1055770). The 2019-nCoV RUO Kit (IDT, #10006713) was used to measure SARS-CoV-2 viral RNA. Viral RNA copy numbers were determined using a standard curve generated with the 2019-nCoV_N_Positive Control (IDT, #10006625). Plaque assays were also performed to further validate the potential existence of live SARS-CoV-2 virus in all samples, before clearance from ABSL-3.

### Behavioral tests

All behavioral tests were conducted in between 15 and 30 dpi in ABSL-2 facility. Mice were transferred to a designated room, 30-minute before behavioral testing. All behavioral tests were performed during the daylight cycle.

#### Foot fault test

The foot-fault test was conducted as previously described. Mice were placed on hexagonal grids of varying sizes and were required to move across the grid, placing their paws on the wires. Each time a paw slipped or fell between the wires during a weight-bearing step, it was counted as a foot fault. The total number of steps taken by each forelimb to traverse the grid was recorded, along with the corresponding number of foot faults for each forelimb.

#### Rotarod test

Test sessions comprised six trials conducted at a variable speed. The protocol began with an initial velocity of 5 rpm for the first 10 seconds, followed by a gradual increase from 5 to 10 rpm over the next 30 seconds, and then a further gradual increase from 10 to 20 rpm between 40 and 80 seconds. The final score was calculated as the average time the mouse remained on the rod across all six trials.

#### Tail suspension test (TST)

During the TST, the mouse was suspended by its tail using tape, with one end anchored to a horizontal bar located 40 cm above the ground. Over the 6-minute duration of the experiment, the mouse shifted from active struggling to a progressively immobile state. The experiment was captured on video, and the duration of immobility was assessed through blinded scoring of the recorded footage after the testing was finished.

#### Open field test (OFT)

Mice were individually placed in a white behavior test box (60 cm × 60 cm × 30 cm, length × width × height) divided into a center field (center, 30 × 30 cm) and a periphery field, where they could freely explore the surroundings for 20 minutes. During each trial, the mouse was positioned in the periphery field, and its movements were digitally recorded by a video camera. Time spent in the center was automatically scored by the customized software.

### Tissue collection and histological analysis

After euthanasia, the brain, liver, and lung tissues were collected and fixed in 4% paraformaldehyde (PFA) overnight. The tissues were then embedded in paraffin and sectioned at 8-µm thickness at the USC Translational Pathology Core. For histological analysis, paraffin sections were deparaffinized in xylene, rehydrated in a series of graded alcohol solutions.

#### Toluidine Blue Staining

To evaluate mast cell activation, we stained sections of lung with 0.1% toluidine blue (Sigma, Cat. #89640). Histological evidence of mast cell activation was detected by the presence of mast cell with dark blue granules.

#### Immunohistochemistry

deparaffinized slides were further subjected to heat-induced antigen retrieval (Vector Laboratories, H-3300). Next, slides were washed with phosphate-buffered saline (PBS, pH 7.4) and blocked with 5% normal donkey serum in PBST (0.2% Triton-X 100 in PBS) for 30 minutes at room temperature. The tissue sections were then incubated with indicated primary antibodies at 4°C overnight, followed by washes with PBST and incubation with secondary antibodies for 1 hour at room temperature. Hoechst 33342 (Thermo Scientific, 62249) was used to label the nuclei. The sections were mounted by ProLong™ Diamond Antifade Mountant (Invitrogen, P36970).

Primary antibodies used in this study include: IBA1 (1:800, FujiFilm CDI R&D Sys, 019-19741), CD68 (1:800, ThermoFisher Scientific, MA5-13324), CD206 (1:200, ThermoFisher Scientific, MA5-32498), and Prealbumin/transthyretin (1:800, proteintech, 11891-1-AP). Secondary antibodies used in this study include: Alexa Fluor 568 donkey anti-rabbit (A10042), Alexa Fluor 568 donkey anti-rat IgG (H+L) (A78946), Alexa Fluor 488 donkey anti-rabbit IgG (H+L) (A21206), Alexa Fluor 488 donkey anti-mouse IgG (H+L) (A21202), and Alexa Fluor 568 donkey anti-mouse IgG (H+L) (A10037). All sections were scanned with Nikon Ti2 confocal microscope equipped with an automated stage; all images were analyzed with ImageJ software.

### Statistical analysis

Data were presented as mean ± standard deviation (s.d.). Statistical analyses were performed with One-way ANOVA or two-tailed unpaired Student’s t-test. *P < 0.05, **P < 0.01, ***P < 0.001 and ****P < 0.0001. NS indicates not significant (P >0.05). P < 0.05 was considered statistically significant. All statistical tests were performed with GraphPad Prism (GraphPad Software).

---

**Supplementary Figure 1.**
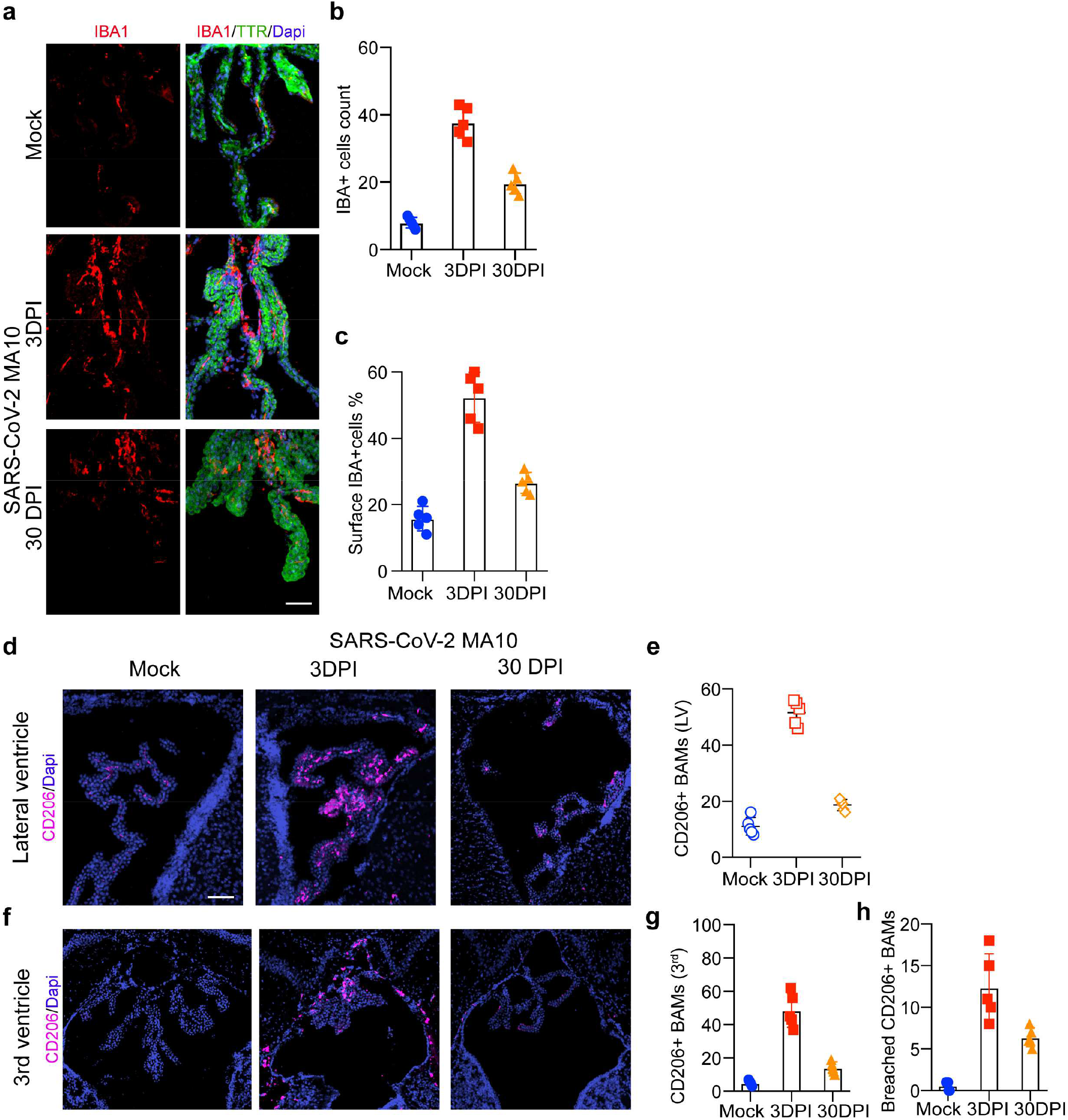
SARS-CoV-2 infection induces accumulation of macrophages at the ChP. **a**. Representative images showing immunostaining for ChP marker TTR and macrophage marker IBA1 in 3^rd^ ventricle of mice with or without SARS-CoV-2 MA10 infection. **b-c**. Quantifications of total ChP macrophage counts (**b**) and surface macrophage counts (**c**) in mock or SARS-CoV-2 MA10 infected mice. **d-e**. Representative images showing CD206-positive cells in the lateral ventricle (LV, **d**) and quantification of CD206-positive BAM counts in LV (**e**) from mock or SARS-CoV-2 MA10 infected mice. **f-h**. Representative images showing CD206-positive cells in the 3^rd^ ventricle (**f**), and quantification of CD206-positive BAM counts in the 3^rd^ ventricle (**g**) and number of CD206-positive BAMs breached from ChP into the ependymal cell layer (**h**) in mock or SARS-CoV-2 MA10 infected mice. Bar = 25 μm. n = 4-5 mice per group.

## References

1. Davis, H. E., McCorkell, L., Vogel, J. M. & Topol, E. J. Long COVID: major findings, mechanisms and recommendations. Nat Rev Microbiol 21, 133–146 (2023).

2. Al-Aly, Z. et al. Long COVID science, research and policy. Nat Med 30, 2148–2164 (2024).

3. Greenhalgh, T., Sivan, M., Perlowski, A. & Nikolich, J.Ž. Long COVID: a clinical update. The Lancet 404, 707–724 (2024).

4. Su, S. et al. Epidemiology, clinical presentation, pathophysiology, and management of long COVID: an update. Mol Psychiatry 2023) doi:10.1038/s41380-023-02171-3.

5. Varatharaj, A. et al. Neurological and neuropsychiatric complications of COVID-19 in 153 patients: a UK-wide surveillance study. The Lancet Psychiatry 7, 875–882 (2020).

6. Bass, D. I. et al. The impact of the COVID-19 pandemic on cerebrovascular disease. Seminars in Vascular Surgery 34, 20–27 (2021).

7. Fraiman, P., Godeiro Junior, C., Moro, E., Cavallieri, F. & Zedde, M. COVID-19 and Cerebrovascular Diseases: A Systematic Review and Perspectives for Stroke Management. Front. Neurol. 11, 574694 (2020).

8. Li, Y. et al. Acute cerebrovascular disease following COVID-19: a single center, retrospective, observational study. Stroke Vasc Neurol 5, 279–284 (2020).

9. Tsivgoulis, G. et al. COVID-19 and cerebrovascular diseases: a comprehensive overview. Ther Adv Neurol Disord 13, 175628642097800 (2020).

10. De Michele, M. et al. Cerebrovascular Complications of COVID-19 and COVID-19 Vaccination. Circ Res 130, 1187–1203 (2022).

11. Munipalli, B., Seim, L., Dawson, N. L., Knight, D. & Dabrh, A. M. A. Post-acute sequelae of COVID-19 (PASC): a meta-narrative review of pathophysiology, prevalence, and management. SN Compr. Clin. Med. 4, 90 (2022).

12. Pretorius, E. et al. Persistent clotting protein pathology in Long COVID/Post-Acute Sequelae of COVID-19 (PASC) is accompanied by increased levels of antiplasmin. Cardiovasc Diabetol 20, 172 (2021).

13. Thaweethai, T. et al. Development of a Definition of Postacute Sequelae of SARS-CoV-2 Infection. JAMA 2023) doi:10.1001/jama.2023.8823.

14. Sherif, Z. A. et al. Pathogenic mechanisms of post-acute sequelae of SARS-CoV-2 infection (PASC). eLife 12, e86002 (2023).

15. Reese, J. T. et al. Generalisable long COVID subtypes: findings from the NIH N3C and RECOVER programmes. eBioMedicine 87, 104413 (2023).

16. Peter, R. S. et al. Post-acute sequelae of covid-19 six to 12 months after infection: population based study. BMJ e071050 (2022) doi:10.1136/bmj-2022-071050.

17. Groff, D. et al. Short-term and Long-term Rates of Postacute Sequelae of SARS-CoV-2 Infection: A Systematic Review. JAMA Netw Open 4, e2128568 (2021).

18. Nath, A. Neurologic complications of coronavirus infections. Neurology 94, 809–810 (2020).

19. Shehata, G. A. et al. Neurological Complications of COVID-19: Underlying Mechanisms and Management. IJMS 22, 4081 (2021).

20. Stefanou, M.-I. et al. Neurological manifestations of long-COVID syndrome: a narrative review. Therapeutic Advances in Chronic Disease 13, 204062232210768 (2022).

21. Filatov, A., Sharma, P., Hindi, F. & Espinosa, P. S. Neurological Complications of Coronavirus Disease (COVID-19): Encephalopathy. Cureus 2020) doi:10.7759/cureus.7352.

22. Spudich, S. & Nath, A. Nervous system consequences of COVID-19. Science 375, 267–269 (2022).

23. Sullivan, B. N. & Fischer, T. Age-Associated Neurological Complications of COVID-19: A Systematic Review and Meta-Analysis. Front. Aging Neurosci. 13, 653694 (2021).

24. Shou, S. et al. Animal Models for COVID-19: Hamsters, Mouse, Ferret, Mink, Tree Shrew, and Non-human Primates. Front. Microbiol. 12, 626553 (2021).

25. Leist, S. R. et al. A Mouse-Adapted SARS-CoV-2 Induces Acute Lung Injury and Mortality in Standard Laboratory Mice. Cell 183, 1070-1085.e12 (2020).

26. Huang, K. et al. Q493K and Q498H substitutions in Spike promote adaptation of SARS-CoV-2 in mice. EBioMedicine 67, 103381 (2021).

27. Wong, L.-Y. R. et al. Eicosanoid signalling blockade protects middle-aged mice from severe COVID-19. Nature 605, 146–151 (2022).

28. Chen, Q. et al. Comparative characterization of SARS-CoV-2 variants of concern and mouse-adapted strains in mice. Journal of Medical Virology 94, 3223–3232 (2022).

29. Huante, M. B. et al. C-type lectin receptor MGL-1 in SARS-CoV-2 disease pathogenesis. The Journal of Immunology 208, 50.30-50.30 (2022).

30. Muruato, A. et al. Mouse-adapted SARS-CoV-2 protects animals from lethal SARS-CoV challenge. PLoS Biol 19, e3001284 (2021).

31. Fernández-Castañeda, A. et al. Mild respiratory COVID can cause multi-lineage neural cell and myelin dysregulation. Cell 185, 2452-2468.e16 (2022).

32. Cui, L. et al. Innate immune cell activation causes lung fibrosis in a humanized model of long COVID. Proc. Natl. Acad. Sci. U.S.A. 120, e2217199120 (2023).

33. Gressett, T. E. et al. Mouse Adapted SARS-CoV-2 Model Induces “Long-COVID” Neuropathology in BALB/c Mice. http://biorxiv.org/lookup/doi/10.1101/2023.03.18.533204 (2023) xdoi:10.1101/2023.03.18.533204.

34. Sefik, E. et al. A humanized mouse model of chronic COVID-19. Nat Biotechnol 40, 906–920 (2022).

35. Peluso, M. J. et al. SARS-CoV-2 and Mitochondrial Proteins in Neural-Derived Exosomes of COVID-19. Annals of Neurology 91, 772–781 (2022).

36. Weinstock, L. B. et al. Mast cell activation symptoms are prevalent in Long-COVID. International Journal of Infectious Diseases 112, 217–226 (2021).

37. Yang, A. C. et al. Dysregulation of brain and choroid plexus cell types in severe COVID-19. Nature 595, 565–571 (2021).

38. Cui, J., Xu, H. & Lehtinen, M. K. Macrophages on the margin: choroid plexus immune responses. Trends Neurosci 44, 864–875 (2021).

39. Xu, H. et al. The choroid plexus synergizes with immune cells during neuroinflammation. Cell 187, 4946-4963.e17 (2024).

40. Giannakopoulos, S., Park, J., Pak, J., Tallquist, M. D. & Verma, S. Post-COVID pulmonary injury in K18-hACE2 mice shows persistent neutrophils and neutrophil extracellular trap formation. Immunity Inflam & Disease 12, e1343 (2024).

41. Jeon, D. et al. Discovery of a new long COVID mouse model via systemic histopathological comparison of SARS-CoV-2 intranasal and inhalation infection. Biochimica et Biophysica Acta (BBA) - Molecular Basis of Disease 1870, 167347 (2024).

42. Singh, A. et al. A murine model of post-acute neurological sequelae following SARS-CoV-2 variant infection. Front. Immunol. 15, 1384516 (2024).

43. Amruta, N. et al. Mouse Adapted SARS-CoV-2 (MA10) Viral Infection Induces Neuroinflammation in Standard Laboratory Mice. Viruses 15, 114 (2022).

44. Lu, H. et al. Potent NKT cell ligands overcome SARS-CoV-2 immune evasion to mitigate viral pathogenesis in mouse models. PLoS Pathog 19, e1011240 (2023).

45. Qiao, H. et al. SARS-COV-2 Induces Blood-Brain Barrier and Choroid Plexus Barrier Impairments and Vascular Inflammation in Mice. http://biorxiv.org/lookup/doi/10.1101/2024.02.09.579589 (2024) xdoi:10.1101/2024.02.09.579589.

